# Plasmacytoid dendritic cells (pDCs) display surface but not membrane-bound IgE across a broad range of total serum IgE levels

**DOI:** 10.1101/2023.11.30.569292

**Authors:** Jonathan H. Chen, Donald W. MacGlashan, Amanda McCormack, John T. Schroeder, Jody R. Tversky

**Affiliations:** Department of Internal Medicine, Lankenau Medical Center, Main Line Health System, Wynnewood, PA, USA; Bloomberg School of Public Health, Johns Hopkins University, Baltimore, MD, USA; Division of Allergy and Clinical Immunology, Department of Medicine, Johns Hopkins University School of Medicine, Baltimore, MD, USA

## Abstract

**Introduction:** Plasmacytoid dendritic cells (pDCs) are unique antigen presenting cells that may be implicated in allergic disease because they bind IgE on their surface and modulate important Th1/Th2 cytokine responses. While conducting in vitro experiments using excess omalizumab to capture IgE, we discovered that this immunoglobulin is not readily removed from pDC to the same extent observed for basophils suggesting that a portion of the IgE on pDC is membrane bound.

**Methods:** Basophils and pDC were prepared from leukopacks using established protocols. In order to isolate PBMCs, blood was also drawn from consenting donors with a wide range in total serum IgE levels. B cells and pDCs were identified by flow cytometry by gating on CD19/CD27 and BDCA2/CD123, respectively. Quilizumab, a mouse anti-human monoclonal IgG1 antibody with specificity only for membrane-bound IgE, was used to detect this molecule in both cell types.

**Results:** When used in vitro at 1.5 mg/ml, omalizumab removed ∼80-90% of the IgE expressed by basophils. In contrast, IgE expression decreased only 30-40% on pDC treated likewise. Upon analyzing pDC for membrane-bound IgE, there was no significant difference in number of target-bound cells and mean fluorescence between the mouse isotype control and quilizumab among pDCs (*P* = 0.125, 0.165). However, the ratio of the proportion of target-bound CD19^+^27^+^ B cells compared to other cells was 32:1 for isotype (*P* < 0.001) and 54:1 for quilizumab (*P* = 0.015).

**Conclusion:** Overall, this study demonstrates that while pDCs express significant levels of FcεRI that binds IgE to the surface, there is no appreciable amount of membrane-bound IgE noted. The reason for omalizumab’s poor ability to remove IgE from pDCs remains unknown.

## Introduction

Plasmacytoid dendritic cells (pDCs) function as weak antigen presenting cells (APCs) but have also been shown to express functional receptors for key elements of both the innate and adaptive immune response.^1,2^ pDCs have the ability to induce proliferation of naïve T cells towards Th1, Th2, or Th17 immune responses and have been implicated in allergic disease process.^1-5^ pDCs express a unique variant of the FcεRI high affinity IgE receptor. Unlike mast cells and basophils, which express prodigious amounts FcεRI on their surface, pDCs express approximately 10-fold less and the receptor itself lacks a beta subunit.^6-8^ It not entirely clear what the specific function of the IgE receptor may be on these cells although it has been implicated to have a role in asthma, atopic dermatitis, allergic rhinitis and other atopic disease.^9-12^ It has been proposed that FcεRI-expressing pDCs use IgE for antigen sampling and/or antigen focusing and modulation.^13^ It is known that pDCs can prime Th1-like responses with secretion of IFN-α and IFN-β.^14^ We have shown that TLR9 expression and activation enhanced these processes while IgE cross-linking inhibits these responses along what may be an important innate and adaptive immune axis.^1,15^ Others later reported similar findings by evaluating the function of TLR7 on pDCs indicating that critical virus-induced IFN-α responses of pDCs can be deficient in patients with allergic asthma.^16,17^

Current literature is mixed on the specific role of pDC-derived IgE signals.^18^ pDCs express a unique trimeric isoform of FcεRI, a key element of the IgE network that is formed by the IgE-binding alpha chain and a dimer of the common Fc receptor gamma chain. It does not express the beta chain, which mast cells and basophils have in the tetrameric isoform, responsible for cell signal amplification.^16^ Increased expression of FcεRI on pDCs correlates with *decreased* severity of food allergy and asthma, suggesting a role of IgE/FcεRI signals in modulating atopic disease. Furthermore, since TLR9 activation of pDCs attenuates IgE/FcεRI mediated production of pro-inflammatory cytokines, suggesting a possible role of the pDC in restraining allergic inflammation.^16,17^ In contrast, allergen immunotherapy increases pDC TLR9-mediated innate immune function, which is impaired at baseline in allergic subjects.^15^

Given the possible role of pDC IgE/FcεRI in allergic inflammation, we aimed to further uncover how IgE may be organized on this cell type. Our preliminary observations suggested that, despite the fact that pDCs display less IgE on their surface, use of the anti-IgE monoclonal antibody, omalizumab only removed a small portion of IgE from the cell surface. This prompted the hypothesis that pDC may secure a significant amount of IgE bound within the cell membrane. The presence of membrane-bound IgE (mIgE) on pDCs is not currently reported or known. We therefore, first more rigorously attempted to remove IgE bound to FcεRI on the surface of pDC in a dose response fashion to compare this outcome with methods previously utilized for basophils and mast cells.^6,19^ We also measured FcεRI expression on the surface of pDCs and correlated this with patient specific total serum IgE levels. We then used quilizumab (a mouse anti-human monoclonal IgG1 antibody) to detect the presence of IgE on the membrane of circulating pDCs and as a control also B cells since these cells are already known to express small amount of membrane bound IgE. Quilizumab is a humanized monoclonal IgG1 antibody directed against the M1’ segment exclusively expressed only on human membrane bound but not receptor bound IgE. We utilized CD19 and CD27 fluorescent antibodies to focus on a juvenile IgE-switched B cell subset in order to increase the probability of identifying membrane bound IgE on these cells in the circulation. The presence of membrane bound IgE (mIgE) on pDC is not currently reported or known. If mIgE is indeed present on pDCs this may help explain why omalizumab only weakly removes IgE from these cells and may suggest an additional role for pDCs in allergic response.

## Methods

### Subject Selection

Blood was drawn from adult volunteers at the Johns Hopkins Allergy and Asthma Center, Baltimore, Maryland according to a protocol approved by the Johns Hopkins Institutional Review Board (IRB). An attempt was made to recruit subjects with a broad range of total serum IgE levels.

### Cell preparation

Blood specimens were processed as described previously.^20,21^ Briefly, whole blood in 10mM EDTA underwent centrifugation at 500g to isolate a buffy coat. Buffy coats were diluted 1:1 (vol/vol) with PAG-EDTA and then subjected to single-Percoll (1.081g/mL) density centrifugation. This produced a PBMC suspension containing mostly mononuclear cells and platelets. Platelets were removed from the PBMC suspension using four low-speed (100g) centrifugations and washes. Cells were fixed in a 1:1 dilution of 2% paraformaldehyde and stored overnight for staining and flow cytometry. To purify basophils and pDCs, a double Percoll gradient was used to generate a basophil-depleted cell (BDC) fraction and an enriched basophil cell (BDC) fraction. Basophils were purified to >99% using negative selection antibodies and beads (STEMCELL Technologies, Cambridge, MA, USA).^22^ The pDC were prepared from the BDC suspension using BDCA-4+ selection after conducting 2 cycles of CD14+ selection to first remove monocytes.^23^

### IgE Stripping of purified pDC and Basophils

Omalizumab (Novartis Pharmaceuticals, East Hanover, NJ, USA) was used to strip IgE from pDCs and basophils (as a control). Omalizumab concentrations used were 0, 100, 500, and 1,500 ug/ml which also included omalizumab stock dilution reagents as a control. We then used a “tube-within-a tube” culture method by inserting a 1.5 ml tube within a 15 ml culture tube with a loosely fit top so that gas exchange is possible. Basophils were cultured at a concentration of 5x10^5 basophils and pDCs at 2.5x10^5 in C-IMDM culture medium which contained IL-3 (10 ng/ml). Cells were cultured for 16h incubation overnight for 16h in a humidified incubator at 37C and 5% CO2. The next day, the cells were pelleted, washed 3x in PBS and then checked for IgE expression using flow cytometry.

### Flow Cytometry

Approximately, 6ml of blood was drawn from each of six consenting individuals, some of whom have self-reported sensitivity to aeroallergens. Peripheral blood mononuclear cells (PBMC) were isolated using standard methods that have been previously described.^20,21^ All cells were first prepared for flow cytometry staining by incubating with a non-specific human IgG blocking antibody (1 mg/ml, MB Biomedicals, Solon, OH, USA). The cells were split into six tubes: two for B cells and four for pDCs. The individual staining conditions are shown below in Table 1. Target antibodies included quilizumab (mouse anti-human IgE, used at 10 µg/ml and obtained courtesy of Genentech, Inc. (San Francisco, CA, USA), anti-human FcεRI (0.5 µg/mL, AER-37, eBioscience/Thermo Fisher, Waltham, MA, USA), and appropriate isotype controls. B cells were gated for using 1:20 dilution commercial mouse anti-human CD19-Alexa488 (BioLegend, San Diego, CA, USA) and 1:20 dilution mouse anti-human CD27-PE (stock 50 ug/mL, BioLegend, San Diego, CA, USA), which has been reported as a marker for juvenile B cells. pDCs were gated for flow cytometry with 1:20 dilution commercial mouse anti-human BDCA-2-Alexa488 (stock 200 ug/mL, BioLegend, San Diego, CA, USA) and 1:20 dilution mouse anti-human CD123 (IL-3)-PE (stock 200 ug/mL, BD Biosciences, Franklin Lakes, NJ, USA). A 1:1,000 dilution of anti-mouse IgG Alexa-647 (stock 1 mg/mL, SouthernBiotech, Birmingham, AL, USA) and was added for secondary labeling of our target antibody. The samples were analyzed to approximately 600,000 total cells via a BD Accuri C6 Plus flow cytometer and mean fluorescence intensity (MFI) was measured.

**Table 1.**
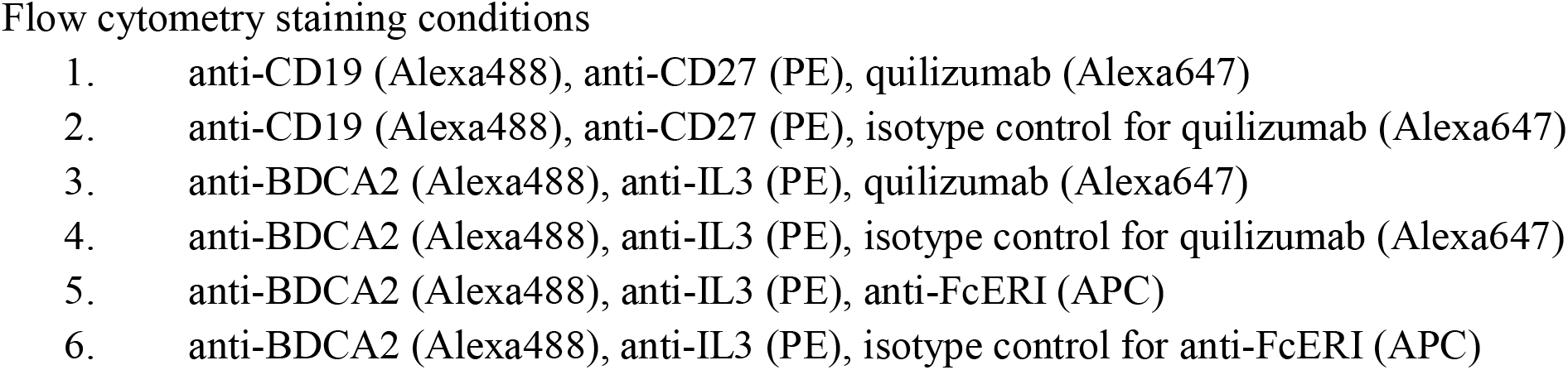

### Statistical Methods

A paired t-test was used to compare proportions of counted cells of both the isotype controls and target antibodies. Data are presented as means and confidence intervals, as appropriate. Statistical analysis was performed with InStat software (GraphPad, San Diego, CA, USA) by a paired t-test or two-tailed non-parametric Mann–Whitney test unless otherwise stated. A linear regression or Spearman’s rank formula was used to calculate correlations as indicated. A *P* value of < 0.05 was considered significant.

## Results

Blood obtained from the six adult subjects had total serum IgE levels ranging from 37-1301 kU/L. As shown in Figure 1A, as a control experiment, omalizumab behaved in a dose-response fashion, with a concentration of 1500 ug/ml effectively removing all IgE from the surface of basophils. In contrast, while pDCs generally have approximately 10-fold less IgE bound to the surface compared to basophils, omalizumab resulted in incomplete removal of IgE (Figure 1B). This suggested that there might be some membrane-bound IgE (mIgE) on the pDC that wouldn’t be removed by omalizumab’s facilitated dissociation characteristic when used at high concentrations. Figure 1C shows the fraction of IgE levels removed from basophils is significantly higher than that of pDCs, which reinforces the finding that IgE is significantly more difficult to strip from pDCs than basophils using omalizumab.

**Figure 1.**
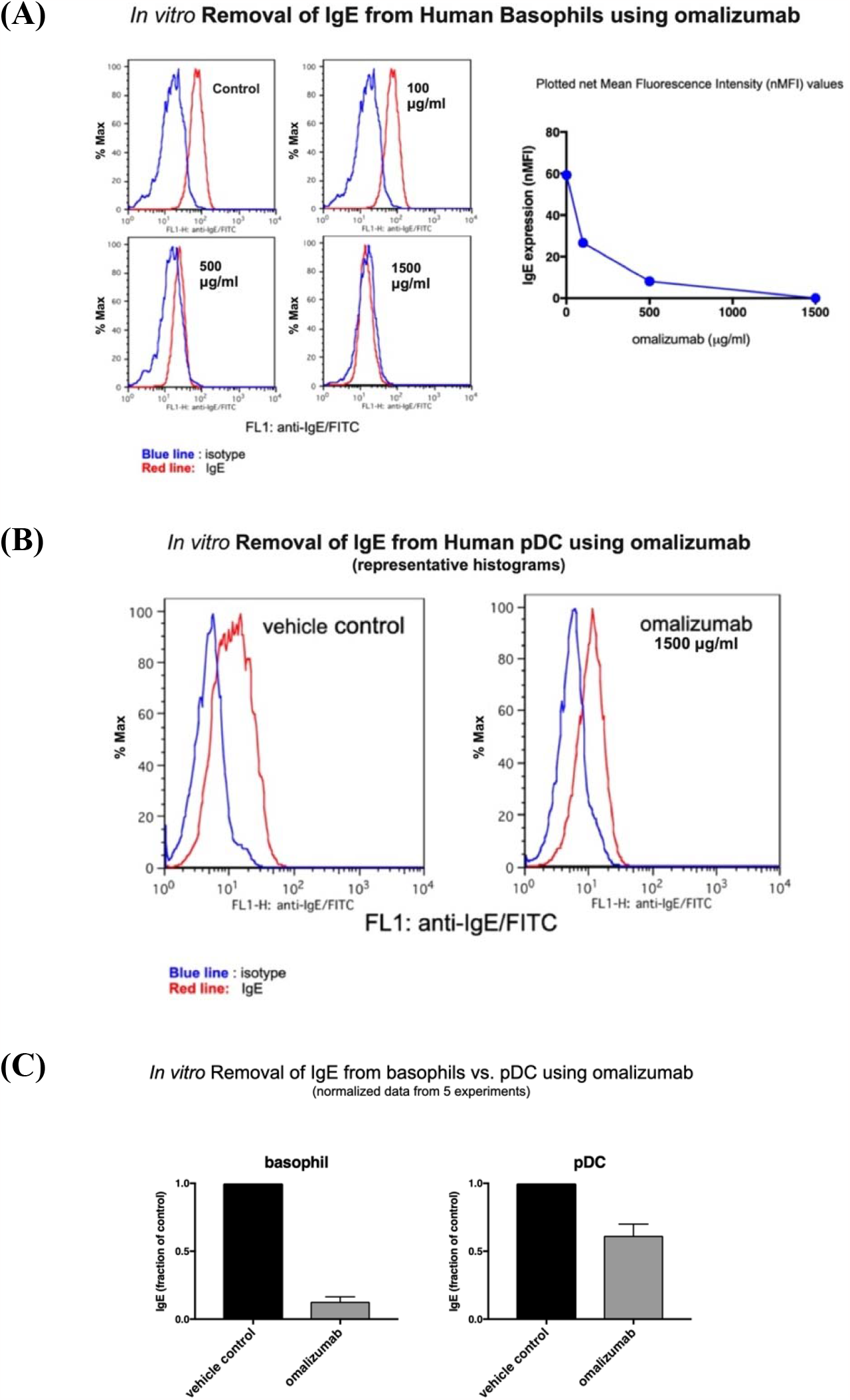
IgE stripping from basophils and pDCs using Omalizumab. (A) Representative histograms of basophils treated with increasing doses of omalizumab are shown. At 500 and 1500 mg/mLs, nearly all of the IgE was removed (*P* <0.0001). (B) Similar histograms of pDCs treated with a 1500 mg/mL high dose of omalizumab are shown. Even at 1500 mg/mL, a large proportion of IgE remained bound to the pDC but did reach significance (*P* = 0.046). (C) Comparing IgE stripping of basophils and pDCs, omalizumab removed significantly more IgE from the basophil compared with the pDC (*P* = 0.02).

The primary outcome was to determine the presence of mIgE on pDCs. As shown in Table 2, BDCA-2+ CD123 (IL-3)+ pDCs did not significantly differ in the proportion of positive cells or MFI between quilizumab and isotype treated cells (0.18% vs 0.19%, *P* = 0.125; and 2,448 vs 2,534, *P* = 0.165, respectively). A representative subject demonstrating the pDC and B cell gating protocols along with quilizumab staining cell selection strategy is shown in Figure 2A-D. The primary outcome was to determine the presence of membrane-bound IgE on pDCs. As shown in Table 2, pDCs identified as BDCA-2+ and CD123+ (IL-3), did not significantly differ in the number of cells or MFI intensity between quilizumab and isotype treated cells 0.18% vs 0.19% (*P* = 0.125) and 2,448 vs 2,534 (*P* = 0.165) respectively. pDCs are uncommon cells in the peripheral bloodstream with typically only about 700 pDC per 500,000 total cells in circulation (0.14%). The number of cells identified as pDC in our study were likewise rare amounting to only about 0.26-0.43% of total PBMC. These data indicate that these rare pDCs do not express an appreciable level of membrane-bound IgE in the bloodstream.

**Table 2.** Quantification of pDCs that bind Quilizumab. BDCA2^+^ IL3^+^ pDCs were isolated in both the isotype and quilizumab-treated cells. The proportion of cells that were positive for Alexa647-isotype or quilizumab did not differ (*P* = 0.125). The MFI levels were similar between the two groups (*P* = 0.165). These data indicate that these rare pDCs do not express an appreciable level of membrane-bound IgE in the bloodstream.

| Subject | Isotype |  |  |  |  | Quilizumab |  |  |  |  | P-value<br>(Target +) | P-value<br>(MFI) |
| --- | --- | --- | --- | --- | --- | --- | --- | --- | --- | --- | --- | --- |
|  | Total #<br>pDCs | % of<br>PBMC | Target<br>(+) | %<br>Target<br>(+) | pDC<br>MFI | Total #<br>pDCs | % of<br>PBMC | Target<br>(+) | %<br>Target<br>(+) | pDC<br>MFI |  |  |
| 1 | 436 | 0.26 | 1 | 0.229 | 2,436 | 699 | 0.41 | 2 | 0.286 | 2,577 | 0.125 | 0.165 |
| 2 | 452 | 0.19 | 1 | 0.221 | 2,461 | 455 | 0.19 | 1 | 0.219 | 2,904 |  |  |
| 3 | 1,426 | 0.26 | 7 | 0.490 | 2,214 | 1,374 | 0.25 | 3 | 0.218 | 1,974 |  |  |
| 4 | 955 | 0.18 | 8 | 0.837 | 2,100 | 891 | 0.17 | 4 | 0.448 | 2,112 |  |  |
| 5 | 1,133 | 0.17 | 2 | 0.176 | 3,100 | 1,162 | 0.18 | 2 | 0.172 | 3,446 |  |  |
| 6 | 669 | 0.09 | 3 | 0.448 | 2,379 | 781 | 0.11 | 2 | 0.256 | 2,189 |  |  |
| **Mean** | **845** | **0.18** | **4** | **0.433** | **2,448** | **894** | **0.19** | **2** | **0.261** | **2,534** |  |  |

**Figure 2.**
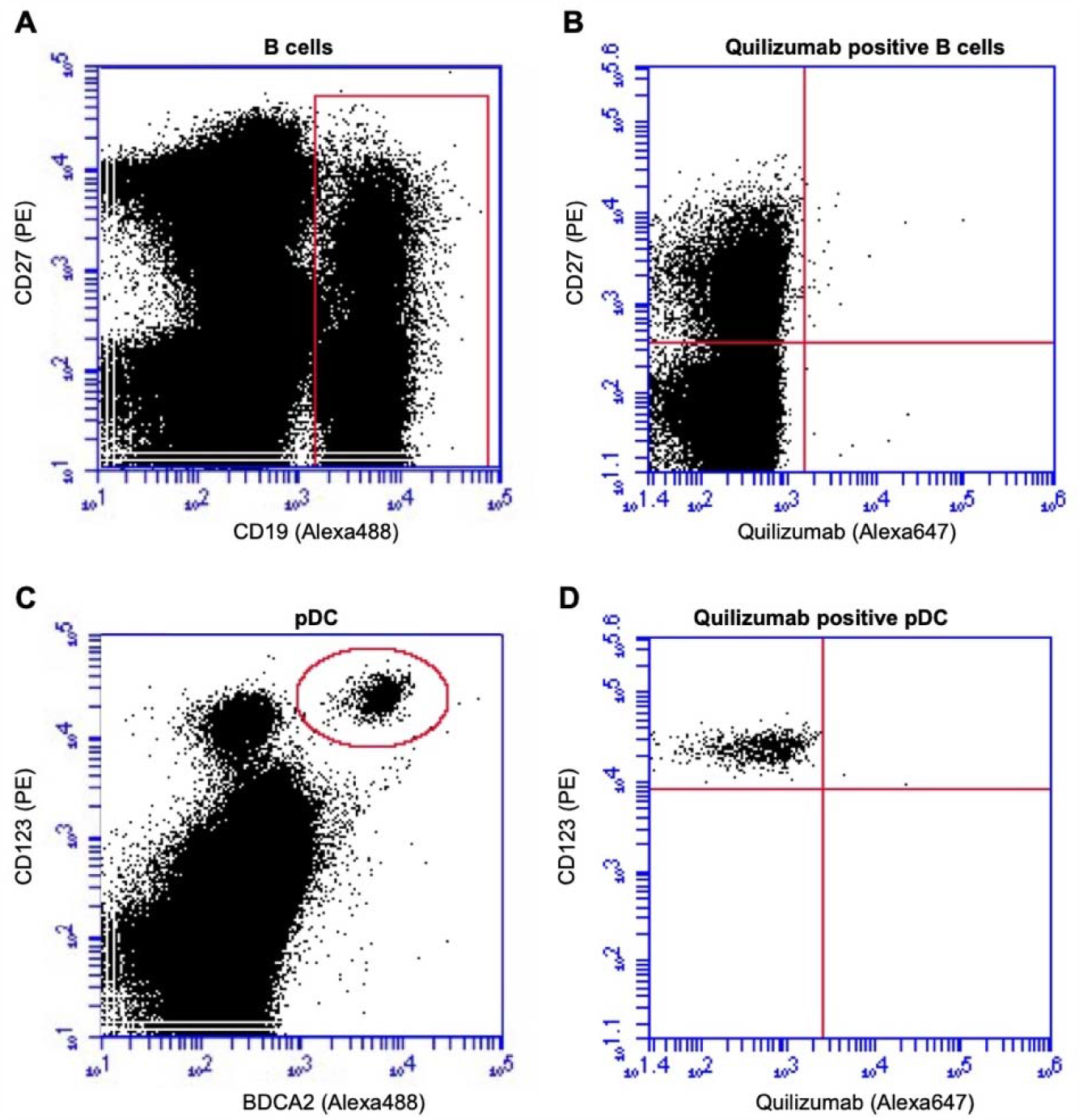
B-cell (A-B) and pDC (C-D) gating strategies. All cells of interest were initially filtered using forward scatter (FSC) and side scatter (SSC) gating based on previously published location of B-cells and pDCs among PBMCs. (A) B cells were then identified with CD19 (Alexa488) and CD27 (PE) indicated by a red rectangular gate. (B) CD19^+^ B cells were then sub-selected for CD27 (PE) and Quilizumab in an effort to identify possible membrane bound IgE bearing B cells (upper right quadrant). In a similar manner (C) pDC were first identified with BDCA2 (Alexa488) and CD123 (PE) indicated by a red circle. (D) Finally, BDCA^+^ pDC were then sub-selected for CD123^+^ (PE) and Quilizumab to identify possible membrane bound IgE bearing pDC (upper right quadrant).

As a control experiment, B cells were separated by cell surface markers to determine whether the membrane-bound IgE target antibody was present in higher proportions in certain subtypes of B cells (Table 3). As expected, the proportion of cells staining positive for quilizumab were highest among the CD19^+^CD27^+^ B cells compared to CD19+CD27-B cells and CD19-sub-groups (0.17%, *P* = 0.008).However, the proportion of cells staining positive for the equivalent isotype-control-stained CD19^+^CD27^+^ B cell sub-group was also highest (0.12%, *P* = 0.014). Overall, among all CD19+ B cells, there was no significant difference in cells staining positive for quilizumab compared to isotype, (*P* = 0.487).

**Table 3.** Quantification of B cell subtypes that bind Quilizumab. Four B-cell subtypes gated on CD19-FITC and CD27-PE were compared. Among all B cells, there was no difference in the proportion of cells binding isotype-Alexa647 and quilizumab-Alexa647 (*P* = 0.487) or MFI (*P* = 0.540). However, for both the isotype- and quilizumab-treated cells, CD19^+^CD27^+^ cells bound significantly more Alexa647 than the other three subtypes (*P* < 0.001, *P* = 0.015).

|  | Isotype |  |  |  | Quilizumab |  |  |  | *P*-value<br>Target + | *P*-value<br>MFI |
| --- | --- | --- | --- | --- | --- | --- | --- | --- | --- | --- |
|  | Total<br>(# cells) | Target +<br>(# cells) | Target +<br>(%) | MFI<br>(mean) | Total<br>(# cells) | Target +<br>(# cells) | Target +<br>(%) | MFI<br>(mean) |  |  |
| CD19-CD27- | 166,354 | 11 | 0.0063 | 5,011 | 169,910 | 6 | 0.0037 | 3,587 | 0.487 | 0.540 |
| CD19+CD27- | 22,969 | 8 | 0.0359 | 1,961 | 25,791 | 8 | 0.0310 | 2,565 |  |  |
| CD19-CD27+ | 213,528 | 13 | 0.0061 | 4,720 | 224,904 | 12 | 0.0053 | 3,818 |  |  |
| CD19+CD27+ | 8,794 | 11 | 0.1232 | 4,096 | 9,012 | 16 | 0.1738 | 3,119 |  |  |

B cells gated with CD19 and CD27 were used as a positive control population because CD19^+^CD27^+^ B cells are thought to more likely express mIgE. Although the total number of CD19^+^CD27^+^ B cells for each patient was greater than the number of pDCs, these CD19^+^CD27^+^ cells are also rare. As shown in Table 4, while there was a trend of higher proportion of CD19^+^CD27^+^ cells staining for quilizumab compared to isotype, this did not quite reach statistical significance (0.174% vs 0.123%, *P* = 0.103). The MFI for isotype vs. quilizumab were also similar among the CD19^+^CD27^+^ B-cells, with the isotype actually having a slightly higher MFI (*P* = 0.043).

**Table 4.** Quantification of CD19^+^CD27^s+^ B-cells that bind Quilizumab. CD19^+^CD27^+^ B-cells were isolated in both isotype- and quilizumab-treated cells. The overall proportion of cells that were positive for isotype-or quilizumab-Alexa647 did not differ (0.12% vs 0.17%, *P* = 0.103)). The overall MFI levels were slightly higher in the isotype group (4,096 vs 3,119, *P* = 0.043). These data indicate that for these developing CD19^+^27^+^ cells, there is increased quilizumab binding, but the rarity of the events make it difficult to distinguish statistically from non-specific binding of the isotype.

| Subject | Isotype |  |  |  | Quilizumab |  |  |  | *P*-value<br>% Target<br>+ | *P*-value<br>MFI |
| --- | --- | --- | --- | --- | --- | --- | --- | --- | --- | --- |
|  | CD19+27+<br>(# cells) | Target +<br>(# cells) | Target +<br>(%) | MFI<br>(mean) | CD19+27+<br>(# cells) | Target +<br>(# cells) | Target +<br>(%) | MFI<br>(mean) |  |  |
| 1 | 1,959 | 4 | 0.2042 | 4,854 | 1,731 | 9 | 0.5199 | 3,373 | 0.103 | 0.043 |
| 2 | 4,886 | 9 | 0.1842 | 4,680 | 4,256 | 14 | 0.3289 | 3,699 |  |  |
| 3 | 13,821 | 14 | 0.1013 | 4,365 | 15,070 | 26 | 0.1725 | 2,033 |  |  |
| 4 | 13,576 | 10 | 0.0737 | 4,505 | 13,248 | 11 | 0.0830 | 2,096 |  |  |
| 5 | 8,557 | 12 | 0.1402 | 2,624 | 10,034 | 13 | 0.1296 | 4,247 |  |  |
| 6 | 9,965 | 16 | 0.1606 | 3,549 | 9,733 | 21 | 0.2158 | 3,265 |  |  |
| **Average** | **8,794** | **11** | **0.1232** | **4,096** | **9,012** | **16** | **0.1738** | **3,119** |  |  |

Given the significantly higher affinity of Alexa647 target to CD19^+^CD27^+^ sub-group cells in both the isotype and quilizumab stained groups, and given the expectation that mIgE expressing B cells would be rare, a comparison of those cells to the rest of the PBMC cells (CD19^-^27^-^, CD19^-^27^+^, CD19^+^27^-^) was calculated (Table 5). In both the isotype and quilizumab groups, there was a statistically significant higher proportion of Alexa647 target-bound cells (*P* < 0.001 and *P* = 0.015, respectively), suggesting that these cells appear to have a higher binding affinity than the CD19-subtypes. A ratio of the proportions was calculated for both groups, comparing CD19^+^27^+^ to the rest of the CD19^-^ cells. More than thirty times more CD19^+^CD27^+^ cells stained positive compared to the CD19^-^ PBMCs. In addition, nearly twice as many quilizumab stained CD19^+^CD27^+^ cells were identified compared to isotype 54.19 vs 32.27 respectively (*P* = 0.234, CI (-63.5,19.7]), although this was not statistically significant due to small sample size and large variance. There was no significant difference in the proportions (*P =* 0.308, *P =* 0.091) or ratio *(P =* 0.355) when comparing CD19^+^27^+^ and CD19^-^27^-^ cells.

**Table 5.** Comparison of CD19^+^27^+^ and other subtypes in both isotype and Quilizumab-treated cells. (A) Proportions of isotype Alexa647-binding cells were compared between CD19^+^27^+^ and CD19+27- and CD19-subtypes. The proportion of Alexa647-bound CD19^+^27^+^ cells was significantly greater than CD19-subtypes (*P < 0*.*001*) The same proportion did not significantly differ between CD19+27+ and CD19+27-cells for either the isotype or quilizumab groups (P = 0.308). The average ratio of the proportions was 32.27. (B) Proportions of quilizumab Alexa647-binding cells were compared between CD19^+^27^+^ and CD19-subtypes. The proportion of Alexa647-bound CD19^+^27^+^ cells was significantly greater than CD19-subtypes (*P* = 0.015). The average ratio of the proportions was 54.19, which was 1.68 times that of the isotype, but did not reach statistical significance. The same proportion did not significantly differ between CD19+27+ and CD19+27-cells (*P = 0*.*355)*.

|  | Subject | CD19+27+ (%) | CD19+27- (%) | CD19- PBMC (%) | CD19+CD27+ : CD19+CD27- (Mean Ratio) | CD19+CD27+ : CD19- PBMC (Mean Ratio) |
| --- | --- | --- | --- | --- | --- | --- |
| Isotype | 1 | 0.2042 | 0.2371 | 0.0237 | 4.35 : 1<br>($P = 0.308$) | 32.31 : 1<br>($P < 0.001$) |
|  | 2 | 0.1842 | n/a | 0.0128 |  |  |
|  | 3 | 0.1013 | n/a | 0.0008 |  |  |
|  | 4 | 0.0737 | 0.0340 | 0.0066 |  |  |
|  | 5 | 0.1402 | 0.0211 | 0.0077 |  |  |
|  | 6 | 0.1606 | 0.0208 | 0.0073 |  |  |
| Quilizumab | 1 | 0.5199 | 0.0746 | 0.0086 | 6.12 : 1<br>($P = 0.091$) | 54.19 : 1<br>($P = 0.015$) |
|  | 2 | 0.3289 | n/a | 0.0032 |  |  |
|  | 3 | 0.1725 | n/a | 0.0017 |  |  |
|  | 4 | 0.0830 | 0.0118 | 0.0046 |  |  |
|  | 5 | 0.1296 | 0.0278 | 0.0073 |  |  |
|  | 6 | 0.2158 | 0.0371 | 0.0081 |  |  |
| | | | | *P* value<br>(isotype vs.<br>quilizumab) | $P = 0.355$ | $P = 0.234$ |

## Discussion

This study is the first to examine the presence of membrane-bound IgE in both circulating B cells and plasmacytoid dendritic cells using quilizumab, a monoclonal antibody that specifically targets the M1 segment that is exclusive to membrane-bound immunoglobulin E. Overall, we did not identify a significant number of circulating plasmacytoid dendritic cells that bind quilizumab. Our study demonstrates that circulating pDCs do not express appreciable amounts of membrane-bound IgE, regardless of the patient’s total IgE level. Therefore, the presence, or lack thereof, of membrane bound IgE on pDCs does not explain our finding that omalizumab insufficiently removes IgE from the surface of pDCs compared to basophils.

We attempted to use circulating B cells, specifically the CD19^+^CD27^+^ subset, as a positive control for quilizumab; however, membrane-bound IgE was determined to be scarce among this cell population as well. Of note, the CD19^+^CD27^+^ cells, which are thought to be a maturing B cell subtype did in fact bind to quilizumab in a much higher proportion compared to the CD19^-^ PBMC (*P* = 0.015) and CD27^-^ B cells *(P* = 0.091). However, while CD19^+^CD27^+^ B cells were more likely to bind quilizumab compared to all CD19^-^ PBMC with a proportion of about 50:1, these cells also bound to the isotype control with a proportion of about 30:1 and therefore did not reach significance. Unfortunately, our study may not have been powered enough to detect a difference in these two assays groups because circulating B cells that possess membrane bound IgE are presumably exceedingly rare.

pDCs are involved in the pathogenesis of multiple allergic conditions including atopic dermatitis, asthma, and chronic rhinitis, in addition to their antiviral immunity and modulation of the innate immune axis through production of type 1 interferons. In the lungs of patients with allergic asthma, for example, pDCs mediate tolerance to inhaled antigen through induction of regulatory T cells compared to conventional DCs which promote Th2 sensitization to inhaled antigen after reaching the lymph nodes.^24^ It has been shown, quite interestingly, that the IgE specific receptor (FcεRI) and TLR9 molecules oppose each other on pDCs.^1^ pDCs have TLR9 molecules through which ligation with CpG DNA favors Th1 responses.Additionally, while pDCs stimulate allergen-dependent T-cell proliferation and Th2 production, CpG-activated pDCs inhibited allergen-dependent proliferation in Th2 memory cells and markedly increased IFN-gamma production.^25^ The results of this study demonstrated that pDCs are not only extremely rare cells in the peripheral blood, but likely do not express membrane-bound IgE. Given the role of pDCs as balance mediators in the Th2-mediated allergic response, a probable postulate is that they do so only through the high affinity IgE receptor, FcεRI and not a membrane bound form.

This study has several strengths and limitations. One of the strengths of this study is that it presented a novel protocol that used an antibody specific to membrane-bound-IgE-specific antibody to stain pDCs. Quilizumab has never been used to determine whether pDCs express membrane-bound IgE. One of the potential limitations of the study is the preferential binding of isotype control antibody to CD19^+^CD27^+^ B cells causing high background. From the data, the MFI range of the isotype is close to that of the target antibody. It is possible that there may have been a small number of quilizumab staining B cells that was masked by the high background binding of the isotype control. Interestingly, like quilizumab, the isotype-control antibody seemed to bind CD19^+^CD27^+^ B cells more specifically than other subtypes as well. Selection of another alternative isotype may have circumvented this.

Lack of significant binding of quilizumab to pDC does not help explain why IgE is more difficult to remove from these circulating dendritic cells compared to basophils. While it has certainly been demonstrated that the FcεRI on pDCs are present in much lower abundance compared to conventional DCs, mast cells, and basophils (with a ratio of about 10:1), the receptor is unique in that it lacks the beta-subunit. Omalizumab does not bind IgE already attached to FcεRI on cells^26^; therefore, omalizumab would presumably bind circulating IgE in the same manner and with the same affinity whether or not it dissociated from a basophils or a pDC. Jardetzky and his collaborators have shown that omalizumab has the capability of inducing facilitated dissociation.^27^ The underlying concept is that omalizumab weakly binds to some portion of IgE not specifically engaged in the receptor pocket, a conformation change is then induced in the IgE that weakens its association with the receptor causing it to more readily dissociate from the receptor. When this happens, it effectively captures the IgE to prevent re-association although the concentrations required to induce this behavior are extreme, i.e. 100 to 1000-fold above stoichiometric levels. This process may be dependent on the precise nature of the molecular environment of bound IgE on pDCs. One hypothesis, therefore is that the unique αγ2 trimeric form of the FcεRI receptor on plasmacytoid dendritic cells, binds IgE more tightly, or differently compared to the αβγ2 tetrameric form expressed on basophils and mast cells. Supporting the concept that the trimer form of FceRI is handled differently by the pDCs, studies suggest that glycosylated FcεRI traffics to the cell surface faster in pDCs compared to basophils^28,29^ and could sequester IgE more readily. Further, studies in transgenic mice expressing human FcεRI suggest that DCs and monocytes internalize IgE to a greater extent than basophils and mast cells and that overall, IgE is bound less frequently to DCs on the surface.^28^ How these differences affect binding and dissociation of IgE on these cells is unknown. More detailed kinetic studies of the interaction of IgE with FcεRI on pDCs and its implications on omalizumab binding needs further exploration.

Overall, this study demonstrates that while circulating pDCs express high levels of FcεRI that binds IgE on the surface, there is not an appreciable amount of membrane-bound IgE detected. Additional studies will need to further address the specific tight interaction of IgE with the unique high affinity FcεRI receptor on plasmacytoid dendritic cells.

